# The Hemopurifier removes extracellular vesicles and microRNAs from renal perfusates following controlled oxygenated rewarming of discarded donor kidneys

**DOI:** 10.1101/2024.08.23.609252

**Authors:** Rosalia de Necochea Campion, Miguel Pesqueira, Paul Vallejos, Cameron McCullough, Alessio Bloesch, Steven P. La Rosa

## Abstract

Kidney transplantation is considered the benchmark treatment for end-stage kidney disease patients. Nevertheless, the demand for donor kidneys is continuously on the rise and the scarcity of suitable kidneys poses a significant hindrance for patients and healthcare providers. One approach is to extend the criteria for the use of kidneys from deceased brain death and deceased circulatory death donors. Use of these organs especially from these extended criteria donors is associated with ischemia reperfusion injury (IRI) and resultant delayed graft function (DGF) as well as increased rates of allograft rejection. One approach to try to lessen these complications as well as increase the time the assessment of organ viability is the use of machine perfusion on recovered kidneys. In this study we obtained perfusates from discarded organs that had undergone Controlled Oxygenated Rewarming. Perfusates were analyzed for extracellular vesicles (EVs), dsDNA associated with EVs and microRNAs. These perfusates were then pumped over the Aethlon Hemopurifier, a plasma separator with an affinity resin containing the lectin *Galanthus nivalis* agglutinin (GNA). Following treatment with the Hemopurifier, a diminution in extracellular vesicles, dsDNA associated with EVs and microRNAs was observed. These results support a future study of the Aethlon Hemopurifier as part of a machine perfusion circuit to explore if the device decreases these mediators in a dynamic circuit and is associated with improved function of retrieved kidneys.

## Introduction

Renal transplantation has emerged as the gold standard treatment for end-stage kidney disease, offering a remarkable improvement in both the quality of life and life expectancy of patients^1^. However, there remains a significant shortage of available donor kidneys as well as a discard rate of approximately 25 % for those organs that are donated^2^. An unavoidable consequence of kidney donation from deceased donors is Ischemia Reperfusion Injury (IRI) as a result of renal artery clamping and cold ischemia associated with traditional static cold storage (SCS). Reperfusion injury leads to delayed graft function (DGF), defined as the need for dialysis in the first 7 days post-transplant, in approximately 30% of renal transplant recipients with rates as high as 55% seen with donors after circulatory death^3,4^. Delayed Graft Function is associated with poor future organ dysfunction as well as allograft rejection^4^.

An advance in the field has been the use of ex vivo machine perfusion of recovered kidneys in the place of traditional static cold storage. Hypothermic Machine Perfusion (HMP) is associated with lower rates of DGF compared with standard cold storage (SCS) in the setting of renal transplantation^5^. Consideration has now been given as to whether Controlled Oxygenated Rewarming (COR) with a period of end-normothemic machine perfusion (NMP) may have advantages over HMP. One potential advantage of gradual rewarming would be less mitochondrial damage and resultant ischemia reperfusion injury^6^. A second potential benefit of this approach is to provide additional time to assess organs and determine viability rather than discard for excessive cold ischemia time^7^.

Early studies with Controlled Oxygenated Rewarming with a period of normothermic machine perfusion by Minor and colleagues have been performed demonstrating its’ feasibility^8,9^. A definitive trial with the Minor protocol has not been published to date. The largest clinical trial done to date incorporating a period of normothermic machine perfusion did not demonstrate a benefit compared to static cold storage on DGF rates of recovered kidneys^10^. A few observations that have been made during NMP may explain this lack of benefit. First, release of extracellular vesicles, which have been implicated in ischemia reperfusion injury^11^ have been noted to be released during normothermic renal perfusion^12^. Second, inflammatory cytokine release has been noted during NMP^13^. Finally, endothelial damage and the development of microthrombi have been noted during NMP as well^14^. All of these findings suggest that the addition of a device that removes inflammatory mediators to a COR circuit might be a valuable adjunct to this treatment of retrieved kidneys.

The Aethlon Hemopurifier is one such device that could serve to remove extracellular vesicles and microRNAs released during the course of COR with NMP. The Hemopurifier (HP) is a combination therapeutic device with a hollow fiber 0.2 µm filter, and an affinity resin produced with a lectin derived from the *Galanthus nivalis* plant (the common snowdrop). The Hemopurifier was designed as an extracorporeal device to be used in blood purification. It combines plasma separation, size exclusion and affinity binding of structures containing mannose^15^ Extracellular vesicles have previously been noted to bind the *Galanthus nivalis agglutinin* lectin in the affinity resin of the Hemopurifier^16^. The Hemopurifier has previously been demonstrated to remove extracellular vesicles from cancer patients when spiked into a buffer solution in vitro^17^ and from a patient with severe COVID-19 infection in vivo^15^. Exosomal miRNAs associated with coagulopathy and acute lung injury were also removed during treatment. In the described proof of concept experiment we examined the ability of the Hemopurifier to remove EVs and microRNAs from perfusate samples collected upon completion of Controlled Oxygenated Rewarming of discarded donor kidneys.

## Methods and Materials

### Ethics Statement

Four human kidneys from the United Network for Organ Sharing (UNOS) program, deemed unsuitable for transplantation, were included in this study. Kidneys discarded for transplantation were offered to 34 Lives by non-profit Organ Procurement Organizations (OPOs). Consent to use these donor kidneys for research was acquired by the OPOs in adherence to standards outlined in the Uniform Anatomical Gift Act of the United States.

### Controlled Oxygenated Rewarming with End Normothermic Machine Perfusion

Discarded kidneys, which had been previously placed on preservation protocols (hypothermic machine perfusion preservation or static cold storage preservation) were perfused with a controlled oxygenated rewarming (COR) protocol designed at 34 Lives (West Lafayette, IN) from an adaptation of the Minor protocol^8^. A mixture of a 750 mL of Steen™ solution (XVIVO Perfusion, Gothenburg, Sweden) and 750 mL of Ringer’s Lactate solution was used as the perfusate, after adding 10 mL of 8.4% Sodium Bicarbonate, 7 mL of 10% Calcium Gluconate, 1 g of ampicillin, and 500 mg of ceftriaxone. The initial 5ºC temperature was gradually increased to 10°C, 17°C, 30°C, and 35°C at timepoint intervals of 30, 60, 75, and 90 minutes. The perfusion flow rate was adjusted to achieve incremental targeted arterial pressure from 30-75 mmHg in the final state period at 35°C. An oxygenator was set to maintain a partial pressure of oxygen above 500 mmHg ensuring adequate oxygen supply to the organ during the final perfusion stage at 35ºC. Total near normothermic perfusion time was 2 hours. Subsequentially, the cumulative renal perfusate solution was collected, frozen, and shipped to Aethlon Medical where they were stored at -80°C until thawed for further analysis.

### Hemopurifier GNA Lectin-Affinity Treatment of Renal Perfusates

A 250 mL volume of COR treated renal perfusate was circulated through the HP cartridge circuit at 200 mL/min using a Spectrum Kr21 peristaltic pump system for a total of 24 volume passes (Figure 1). Each HP cartridge contained 40 grams of GNA affinity resin (Aethlon Lot No. RD070523). As shown in Figure 1, during HP treatment the kidney perfusate passes through the packed column of GNA affinity resin promoting molecular interactions between the donor renal perfusate and the resin.

**Figure 1.**
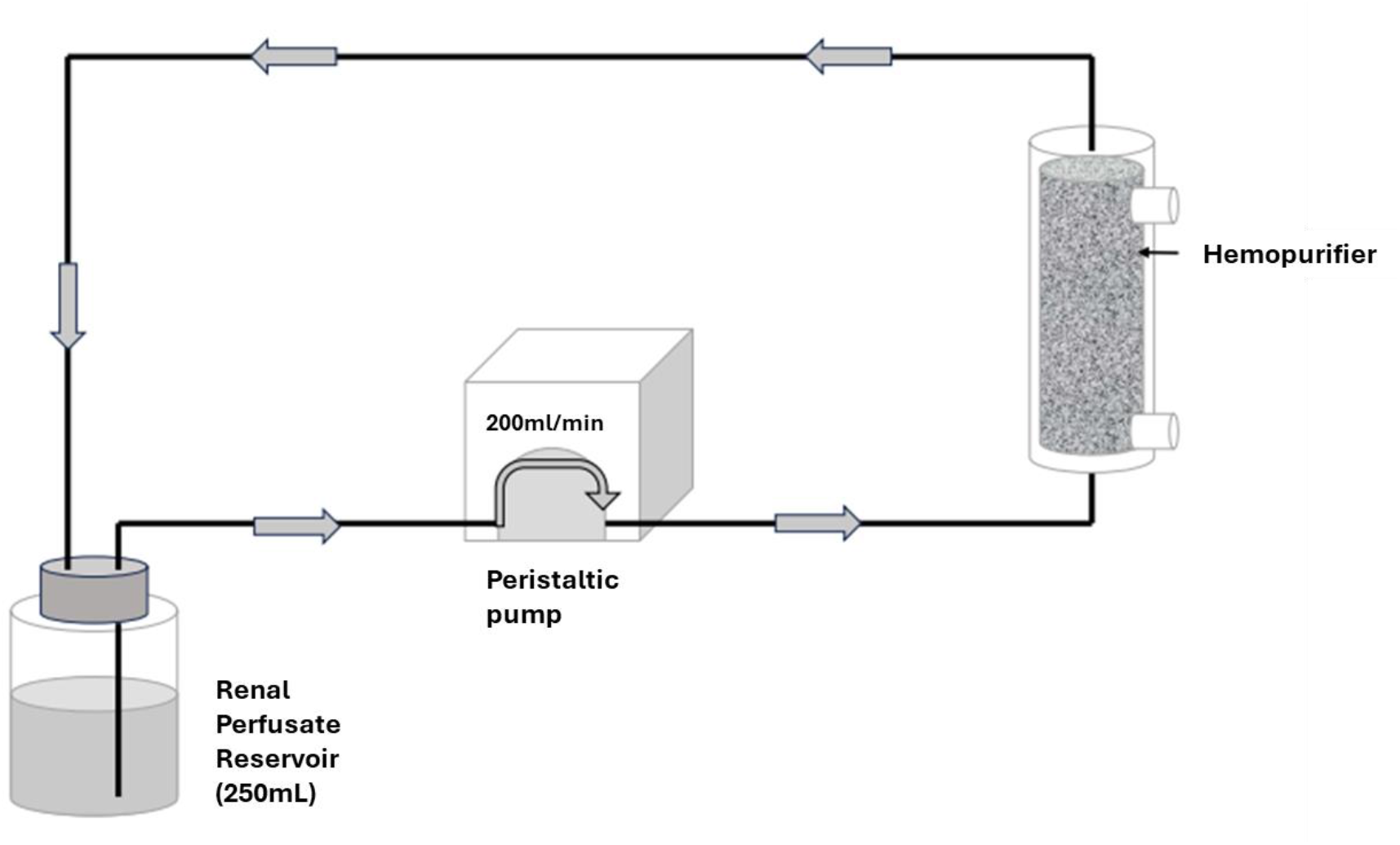
Renal Perfusate Hemopurifier® Circuit Set-Up. This figure depicts the benchtop configuration of the circuit used to treat the kidney perfusates with the Hemopurifier cartridge. Kidney perfusates were circulated through the Hemopurifier where they passed through a packed column of GNA affinity resin for a total of 24 cycles (30 min). Perfusate samples were collected after treatment and compared to pretreatment samples to identify differences in EV and miRNA contents.

### EV quantification

Concentration of EV and dsDNA-associated EV nanoparticles was determined through the use of high sensitivity nanoparticle NanoFCM Flow Analyzer U30 (NanoFCM, Nottingham, UK). Co-staining was performed by first mixing samples directly with 2 μM MemGlow 640 nm (MemGlow™ 640: Fluorogenic Membrane Probe, Cytoskeleton) at a 1:1 ratio for quantification of EV particles. 2 μL of MemGlow stained sample was diluted into 98 μL of reverse osmosis (RO) purified water (Sartorius) and stained with SYTOX Green Ready Flow Reagent (Invitrogen), a membrane impermeable nucleic acid dye. The nucleic dye itself was diluted 10-fold, used to stain at a ratio of 1:1 with the diluted MemGlow stained sample and incubated at room temperature for 15 minutes prior to single event counting analysis on the NanoFCM instrument.

### RNA Isolation

Total RNA was isolated from both pre- and post-HP circulated kidney perfusates. The RNA isolation kit used was the Plasma/Serum RNA purification Mini Kit (Norgen Biotek Corp, Thorold, Canada) according to the manufacturer’s protocol. In brief, 200 µL of kidney perfusate were thoroughly mixed with the lysis buffer and three spike-in exogenous miRNAs were added (ath-miR159a, cel-miR-248, osa-miR414) at a 1.6 pM concentration before processing on the spin column. Purified RNA was eluted with 20 µL of elution buffer and quantified using the Nanodrop One Spectrophotometer (ThermoFisher Scientific).

### NanoString nCounter miRNA Assay

A 10 µL aliquot of purified RNA (∼25 ng/µL) from each sample was provided to the NanoString Services Laboratory at Canopy Biosciences (Canopy Biosciences, Hayward, CA). The samples were prepared for NanoString nCounter miRNA expression profiling according to the manufacturer’s protocol. Analysis of miRNA raw data (RCC data files) was performed using the NanoString nSolver analysis software (version 4.0). This R-based software package is the only tool that allows for data normalization using the NanoString ligation controls, in addition to its other important analytical elements including signal density quality controls, background correction, and differential expression analysis^18^. Before data normalization, background thresholding was performed, setting the threshold count value to 25 reads based on the mean count of the negative controls plus two standard deviations. Data normalization was performed using the geometric mean of the exogenous spike-in miRNA values (ath-miR159a, cel-miR-248, osa-miR414) in addition to the nCounter positive ligation controls. Differential analysis was then performed to assess miRNA targets significantly altered by Hemopurifier Treatment.

### Statistical Analysis

Statistical analyses on extracellular vesicle data sets was performed using GraphPad Prism 8 software. A one-tailed paired t-test was used to assess significant changes in the EV measurements in treated perfusate samples. The accepted level of significance was *P*≤ 0.05 indicated by one asterisk (see figure legends). Statistical analysis of the miRNA datasets was performed using the NanoString nSolver analysis software (version 4.0) to identify significant changes in treated perfusate samples. The nSolver False Discovery Rate (FDR) filter was applied to control for error in multiple hypothesis p-values testing and identify only those changes that were truly significant in this treatment model. Significance was defined as p≤ 0.05.

## Results

### Donor Characteristics

HP treated perfusates were obtained from four different deceased renal donors (2 DCD and 2 DBD) with a median Kidney Donor Patient Index (KDPI) of 75.5%, ranging from moderate to high risk of graft rejection/function^19^. Median Cold ischemia time was 28 hours and 12 minutes and other health and demographic data contributing to the calculated KDPI values is shown in Table 1.

**Table 1.**
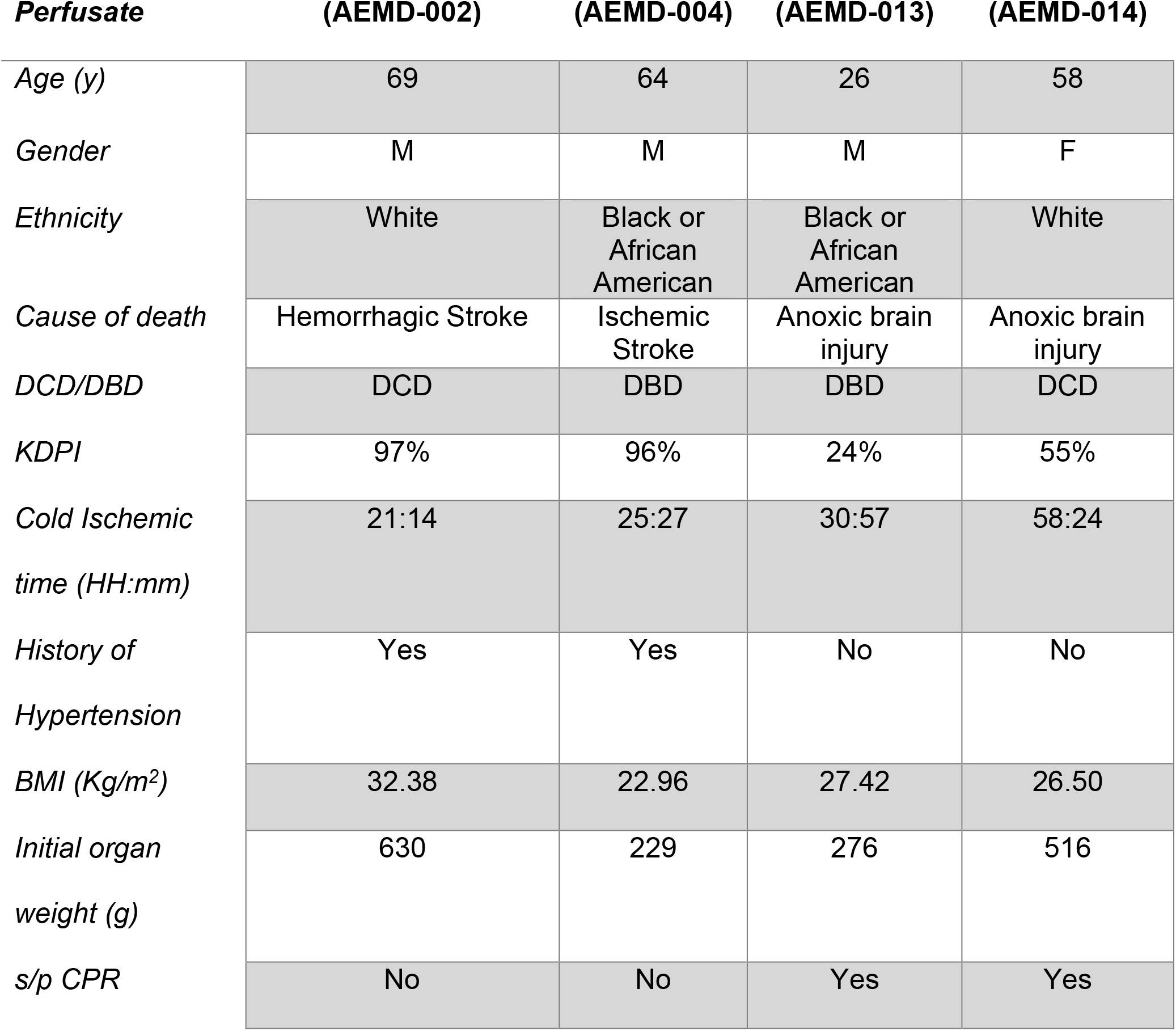
Renal Donor Characteristics and Demographics.

### EVs and a DNA-associated subpopulation of these nanoparticles are depleted by Hemopurifier treatment

Nanoparticle flow cytometry analysis of renal perfusates was performed to quantify the concentrations of small EVs, 40-200 nm size range, and larger EVs in the 100-500 nm range pre- and post-HP treatment. Although small EVs and larger EVs inherently have some size overlap, it is important to distinguish between these subpopulations because they are known to have different cellular origins and distinct cargo with different implications in renal transplant outcomes^20,21^.

As shown in Figure 2a, there is a statistically significant reduction of small EV particles in 3 out of 4 of the HP treated renal perfusates. For example, AEMD-04 had a measured average initial concentration of 1.74E+10 EV particles/mL and a post-HP treatment concentration of 1.48E+9 EV particles/mL, signifying over a 10-fold statistically significant reduction. In contrast, although EV reduction was nearly 4-fold in AEMD-02 treated perfusates, this difference did not quite reach statistical significance (p = 0.052). Comparison of pre- and post-HP small EV particles/mL(40-200 nm size) from AEMD-02, AEMD-04, AEMD-13, and AEMD-14 perfusates demonstrate a 73.1%, 91.5%, 86.0%, and 94.5% vesicle depletion, respectively. Further, particle colocalization quantification analysis for EVs in tandem with dsDNA showed a significant reduction, of similar extent to the EV depletion, of this DNA-associated EV subpopulation across the four renal perfusate samples, as shown in Figure 2b. The small DNA-associated EV particle concentrations of AEMD-02. AEMD-04, AEMD-13, and AEMD-14 perfusates were depleted by 95.2%, 84.9%, 77.7%, and 96.4%, respectively. These reductions were all found to be statistically significant (Figure 2b), except for AEMD-02 where the difference did not quite reach statistical significance (p = 0.058).

**Figure 2.**
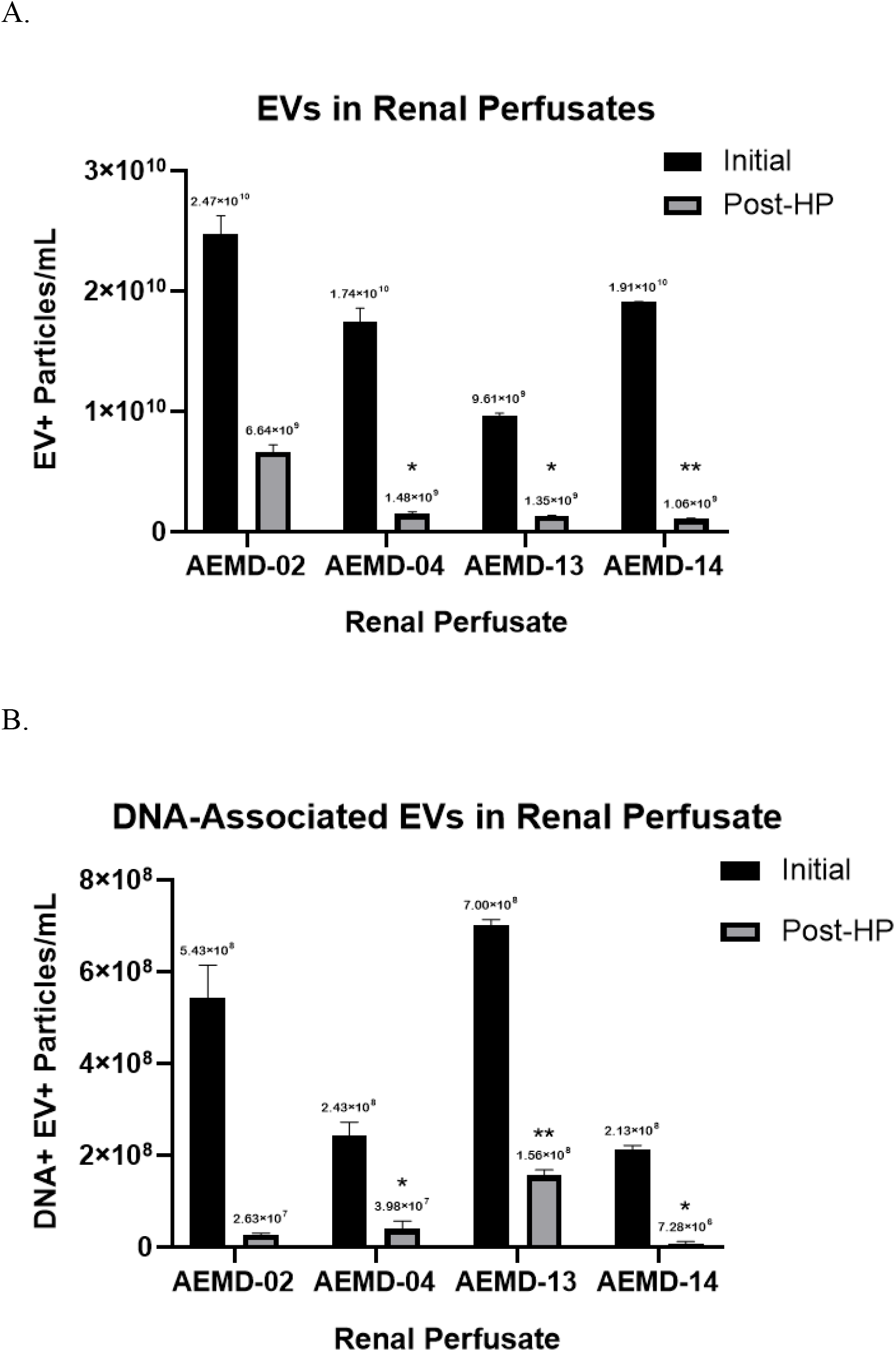
Renal perfusates contain small extracellular vesicles that are removed by Hemopurifier treatment. (A) NanoFCM nanoparticle flow cytometry analysis demonstrates that the Hemopurifier removes the majority of small EV particles (40-200nm nanovesicles) and (B) of dsDNA loaded EV particles in treated renal perfusates. The data shown represents the mean and standard error of 2 technical replicates measured from each perfusate sample.(* p ≤ 0.05; **p ≤ 0.01)

To gain further insight into the HP’s ability to remove EVs from renal perfusates, we adjusted our nanoparticle flow cytometry parameters for the detection of larger EVs ranging from 100-500 nm in diameter size. Although the number of EVs in the 100-500 nm size range in the renal perfusates samples is less abundant than those in the 40-200 nm range, there is also statistically significant reduction of such larger EVs by the HP treatment. As shown in Figure 3a, these larger EV populations were all significantly reduced by 94.6%, 98.9%, 71.0%, and 99.4% in the AEMD-02, AEMD-04, AEMD-13, and AEMD-14 perfusate samples, respectively. Moreover, the subpopulation of DNA-associated larger EVs was also significantly reduced, as shown in Figure 3b, with an average depletion of 98.2%, 100%, 89.7%, and 100% for each of the four renal perfusate samples, respectively.

**Figure 3.**
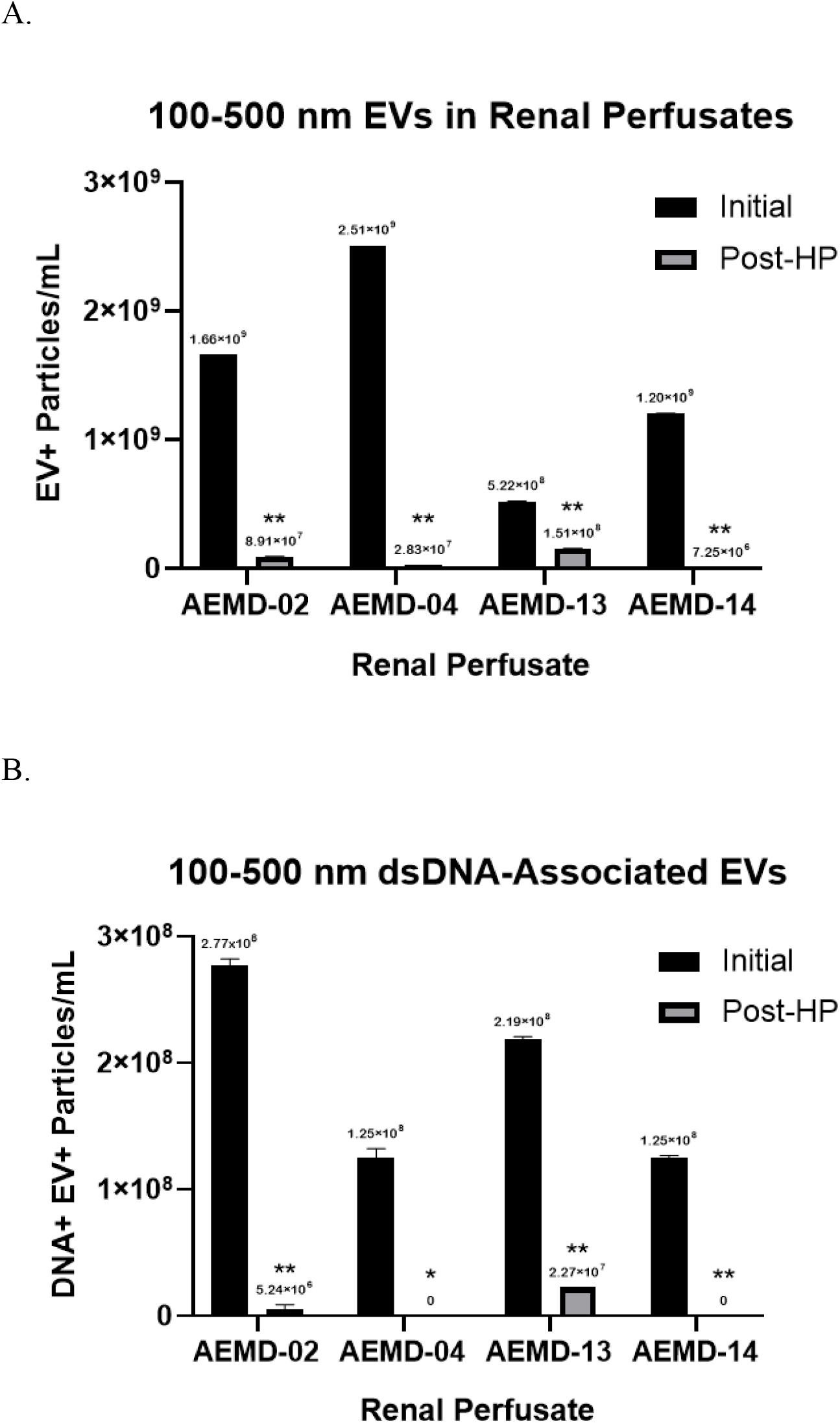
Larger extracellular vesicles within the fiber pore-size filtration capacity of the Hemopurifier are removed from treated renal perfusates. (A) Differences in 100-500 nm EV counts from the four treated renal perfusates. (B) pre- and post-HP concentrations of DNA-associated large EV particles. Measurements for AEMD-04 and AEMD-14 post-HP DNA-associated EV particles were below the instrument’s limit of detection which was 5.22E+6 and 5.58E+6/mL, respectively. Data shown represent the mean and standard error of 2 technical replicates measured from each perfusate sample. (* p ≤ 0.05; **p ≤ 0.01)

### Perfusate miRNAs are depleted by Hemopurifier treatment

Pre- and post-HP treated RNA was submitted for miRNA analysis. Out of the potential 798 miRNAs present on the NanoString Human miRNA Panel, 341 miRNAs were detected in the perfusate with content levels that could be measured above the background threshold. Of those 341 detectable miRNAs, 235 were depleted by HP treatment (data not shown). Initially, 22 of these miRNAs were found to be significantly depleted (p< 0.05) by treatment (Supplemental Table 1). However, when a false discovery rate filter was applied to the data to control for the type I error in our multiple testing model, we found that 5 of these miRNA were identified as significantly reduced by HP treatment (Table 2). The average reduction in this group was 67.8% with species such as hsa-let-7a-5p and hsa-miR-29b-3p demonstrating the highest levels of 86.5% and 77.7% depletion, respectively.

**Table 2.** Significantly depleted miRNA in Hemopurifier treated perfusates.

| Probe Name | Accession # | 24P vs.<br>ctrl | FDR (p-value) | Percent<br>Reduction |
| --- | --- | --- | --- | --- |
| hsa-let-7a-5p | MIMAT0000062 | -7.4 | 0.00 | -86.5 |
| hsa-miR-148b-3p | MIMAT0000759 | -1.78 | 0.04 | -43.7 |
| hsa-miR-148a-3p | MIMAT0000243 | -2.36 | 0.04 | -57.7 |
| hsa-miR-29b-3p | MIMAT0000100 | -4.48 | 0.05 | -77.7 |
| hsa-miR-99a-5p | MIMAT0000097 | -3.75 | 0.05 | -73.3 |

## Discussion

This study has provided significant insights into the release of extracellular vesicles and microRNAs during Controlled Oxygenated Rewarming with end near-Normothermic Machine Perfusion (NMP). Importantly, we observed large quantities of extracellular vesicles, EVs with DNA cargo, and microRNAs released during the procedure. Furthermore, the Aethlon Hemopurifier® removed these potential donor organ transplant mediators from the end perfusate. This data raises the hypothesis that incorporation of the device into the COR treatment of a recovered kidney might improve viability of extended criteria kidneys and important outcomes following kidney transplantation.

Extracellular vesicles or EVs are heterogenous lipid bilayer membrane nanoparticles (20-10,000 nm in diameter) released by all cell types that participate in cell-to-cell communication^22^. Within the EVs are cargo including nucleic acids, proteins, cytokines and non-coding microRNAs involved in regulation of transcription.

Extracellular vesicles released as a consequence of hypoxia have been implicated in ischemia reperfusion injury as well as Delayed Graft Function^11^. Donor derived EVs have also been implicated in the development of allograft rejection^23^. Woud et al is but one group that demonstrated an increase in EV number during the course of normothermic machine perfusion in extended criteria donor discarded kidneys^12^.

Rutman at al examined hypothermic kidney perfusate EVs and found that they were < 500nm in diameter and that they contained donor HLA markers^24^. We were able to demonstrate a 73% to 94% reduction in total EV concentration when renal perfusates were run over the Aethlon Hemopurifier. Furthermore, we were able to demonstrate removal of EVs up to 500nm in size (Supplemental Figure 1). This larger size range is known to include donor microvesicles and apoptotic bodies with distinct molecular cargo which can contribute to graft dysfunction^21^.

The nanoparticle flow cytometry analysis results also demonstrate the Aethlon Hemopurifier®’s ability to remove dsDNA-associated EVs from renal perfusates. The small EVs (40-200 nm in diameter) most likely contain dsDNA molecules associated to their surface due to the membrane-impermeable nature of the Sytox green nucleic acid dye co-localized with the phospholipid bilayer stain. Although the mechanism through which this occurs is not well understood, this phenomenon has been observed in other small EV populations and might be worth exploring further^25^. More importantly though, DNA from EVs can induce the recruitment of immune cells in various conditions by incrementing the expression of type I interferons and cytokines through the activation of the DNA damage receptor cyclic GMP-AMP synthase (cGAS)-stimulator of interferon genes (STING) pathway^25,26^. Therefore, the elimination of dsDNA-associated EVs in renal perfusates by HP treatment suggests another therapeutic mechanism to reduce inflammation and immune response after transplantation.

One specific cargo component of extracellular vesicles that has garnered attention in renal perfusates is microRNAs. Gremmels et al specifically examined renal donor preservation fluid and found 10 kidney perfusate extracellular vesicle (KP-EV) miRNAs that differentiated delayed graft function from immediate graft function^27^. In the study by Rutman he found increased levels of miRNA 218-5p that was associated with delayed graft function as a result of an increased TH17 /T regulatory cell ratio^24^. Separate studies Li and Ding have identified exosomal miRNAs associated with renal ischemia/reperfusion injury respectively that could be blocked by antagonist treatment^28,29^. This data in total raises the possibility that EV miRNAs could be targets for removal during machine perfusion of recovered kidneys.

Our study examined the largest number of microRNAs in perfusates to date following normothermic machine perfusion of discarded kidneys. After controlling for false discovery, we observed a significant reduction in five microRNAs after the perfusates were passed over the Hemopurifier. The Aethlon Hemopurifier® treatment led to the significant depletion of hsa-let-7a-5p, has-miR-148b-3p, has-miR-148a-3p, has-miR29b-3pb and has-miR-99a5p. These data appear to suggest that the mechanism of miRNA removal is affinity based and that miRNA species with significant depletion may be packaged in EVs with specific mannose-rich glycosylation patterns. Let-7a-5p has been found to be a regulator promoting renal dysfunction. Upregulation has been noted in Multiple Myeloma patients^30^. Inhibition of the let-7a/TGFCR1 signaling has ameliorated diabetic nephropathy while let-7a upregulation has been demonstrated to enhance hyperinflammation by increasing cell proliferation and NFkappa-b activation in SLE cell models^31,32^. In prior studies miR148b-3p has been associated with IgA nephropathy^33^. Upregulation of this microRNA has been seen in localized irradiation injury in a mouse model and following LPS injection in a pig model^34,35^. The microRNA 148a-3p has been associated with renal fibrosis in a unilateral ureteral ligation model^36^. Additionally, miR148a-3p has been shown to promote polarization to an inflammatory M1 phenotype in macrophage as well as involved in pro-inflammatory cytokine release^37,38^. The miR 29b-3p has been noted to be upregulated in renal chronic allograft patients but its role may be protective and not injurious^39^. Finally, miR 99a-5p has been associated with urinary EVs in renal ANCA-associated vasculitis but has an unclear role in renal diseases^40^.

Despite the promising findings of this study there are limitations to consider. The study was done with spent renal perfusates following the completion of the COR treatment. As such we did not examine the kinetics of production or removal of these EVs and microRNAs over the course of the Hemopurifier treatment. Additionally, we do not know the effects on the histopathology of the recovered organ or the effects of removal of these mediators on renal function. We also only studied effects on perfusates following a COR protocol while Hypothermic Machine Perfusion is the current standard of care for ex vivo treatment of recovered kidneys. The next step would be to compare mediator removal, renal function and histopathology in a COR circuit performed with or without incorporation of the Hemopurifier on discarded kidneys.

Ultimately, we would need a clinical trial demonstrating that incorporation of the Hemopurifier into renal perfusion improves important clinical endpoints in transplant recipients such as DGF, graft survival or rejection rates to justify its use.

## Abbreviations

cGAS: cyclic GMP-AMP synthase
COR: Controlled Oxygenated Rewarming
DBD: Donation after Brain Death
DCD: Donation after Circulatory Death
DGF: Delayed Graft Function
DNA: Deoxyribonucleic acid
dsDNA: double-stranded DNA
EV: Extracellular Vesicle
HMP: Hypothermic Machine Perfusion
HP: Hemopurifier
IRI: Ischemia Reperfusion Injury
NMP: Normothermic Machine Perfusion
RO: Reverse Osmosis
SCS: Static Cold Storage
STING: Stimulator of Interferon Genes
UNOS: United Network for Organ Sharing

## Acknowledgments

The authors would like to thank 34 Lives for sending their COR renal perfusates to Aethlon Medical for use in this study.

