## Supplemental Figure 1 for "The Hemopurifier removes extracellular vesicles and microRNAs from renal perfusates following controlled oxygenated rewarming of discarded donor kidneys"

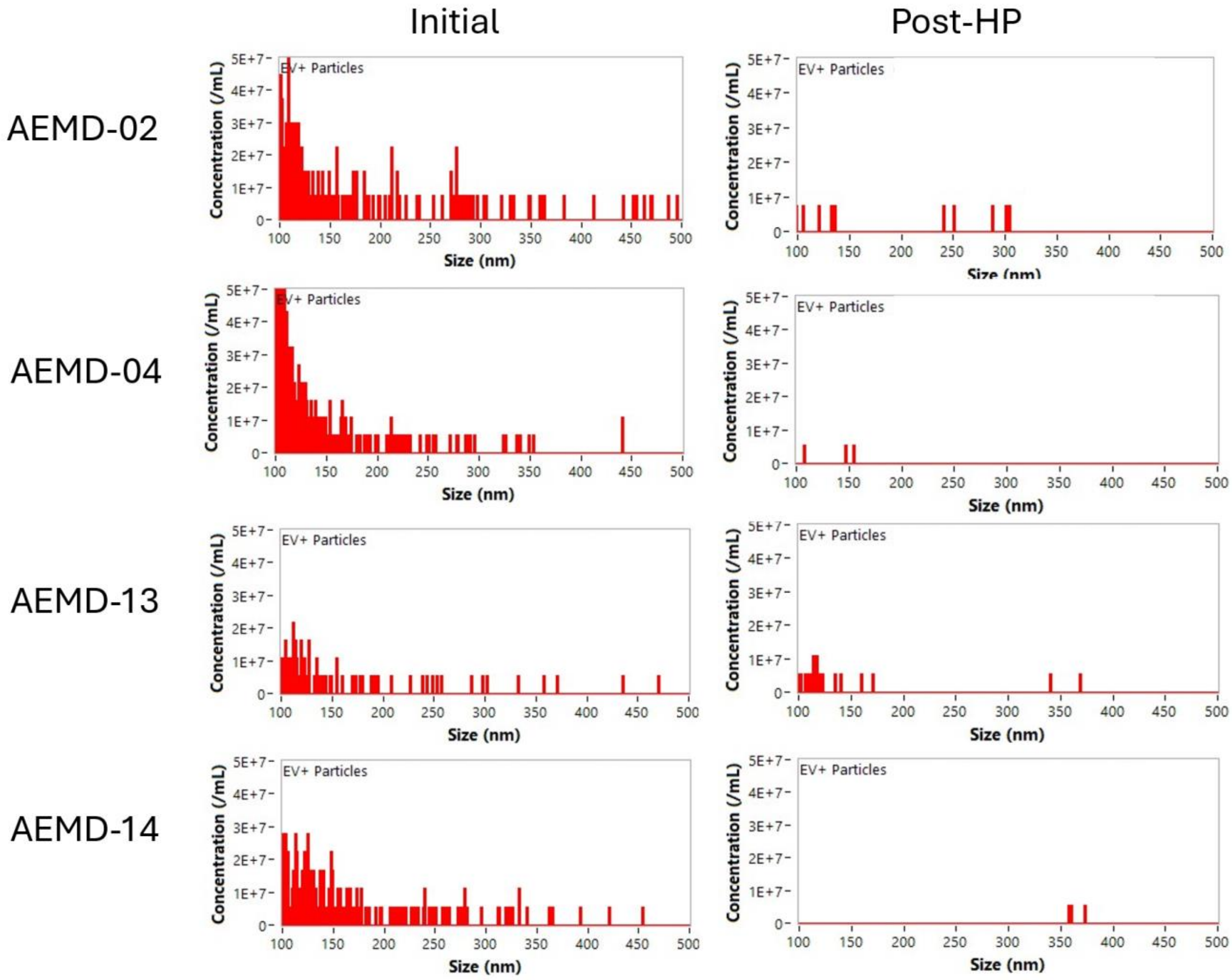


***Supplemental Figure 1. Size distribution graphs for larger EV particles depleted by HP-treatment.*** *The concentration of EV particles per mL at 0.5 nm size intervals, found at the 100-500 nm range, is plotted for the four renal perfusates pre- and post-HP treatment. The distribution data shown was collected from one representative technical replicate of each perfusate sample analyzed in this study.*
