## Supplemental Table 1 for "The Hemopurifier removes extracellular vesicles and microRNAs from renal perfusates following controlled oxygenated rewarming of discarded donor kidneys"

***Supplemental Table 1.*** *Perfusate miRNA that appear to be significantly depleted by Hemopurifier treatment when no False Discovery Rate correction is applied to the data.*

| Probe Name | Accession # | 24P vs ctrl | p-value | Percent Reduction |
|---|---|---|---|---|
| hsa-let-7a-5p | MIMAT0000062 | -7.4 | 0.00 | -86.5 |
| hsa-miR-148b-3p | MIMAT0000759 | -1.78 | 0.00 | -43.7 |
| hsa-miR-148a-3p | MIMAT0000243 | -2.36 | 0.00 | -57.7 |
| hsa-miR-29b-3p | MIMAT0000100 | -4.48 | 0.00 | -77.7 |
| hsa-miR-99a-5p | MIMAT0000097 | -3.75 | 0.00 | -73.3 |
| hsa-miR-25-3p | MIMAT0000081 | -1.98 | 0.01 | -49.5 |
| hsa-miR-15b-5p | MIMAT0000417 | -2.88 | 0.01 | -65.2 |
| hsa-miR-142-3p | MIMAT0000434 | -2.36 | 0.01 | -57.7 |
| hsa-miR-660-5p | MIMAT0003338 | -2.57 | 0.01 | -61.0 |
| hsa-miR-141-3p | MIMAT0000432 | -2.02 | 0.01 | -50.6 |
| hsa-miR-126-3p | MIMAT0000445 | -2.45 | 0.02 | -59.2 |
| hsa-miR-15a-5p | MIMAT0000068 | -2.33 | 0.02 | -57.1 |
| hsa-miR-324-5p | MIMAT0000761 | -2.59 | 0.03 | -61.5 |
| hsa-miR-518b | MIMAT0002844 | -2.44 | 0.03 | -59.0 |
| hsa-miR-4454 | MIMAT0018976 | -4.06 | 0.03 | -75.4 |
| hsa-miR-19b-3p | MIMAT0000074 | -2.33 | 0.03 | -57.1 |
| hsa-miR-199b-5p | MIMAT0000263 | -2.39 | 0.04 | -58.2 |
| hsa-miR-495-3p | MIMAT0002817 | -2.1 | 0.04 | -52.3 |
| hsa-miR-424-5p | MIMAT0001341 | -2.3 | 0.04 | -56.5 |
| hsa-miR-193a-5p | MIMAT0004614 | -2.49 | 0.04 | -59.9 |
| hsa-miR-30d-5p | MIMAT0000245 | -1.57 | 0.05 | -36.2 |
| hsa-miR-377-3p | MIMAT0000730 | -1.84 | 0.05 | -45.6 |
